# Species distribution modeling for conservation science: new predictor layers, reproducible code, and an evaluation of California protected areas

**DOI:** 10.1101/2025.01.23.634559

**Authors:** Zachary G. MacDonald, Joscha Beninde, Kentaro Matsunaga, Bo Zhou, Thomas W. Gillespie, H. Bradley Shaffer

## Abstract

**Aim:** Our study provides foundational resources for future SDMing: methods for generating fine-scale, equal-area predictor datasets and best-practice SDM guidelines. We also provide reproducible code to streamline their implementation.

**Location:** Southwestern North America

**Methods:** Using over 215,000 research-grade iNaturalist occurrence records for 127 species of conservation concern or scientific interest in California and surrounding area, we quantified and compared SDM performance between two predictor datasets that differ in their source of bioclimatic data, spatial resolution, and coordinate reference system: one generated using ClimateNA software (resolution = 300 x 300 m; NAD83/California Albers) and the other using existing WorldClim data (varying resolution = ∼669-797 x 926 m; WGS84). We also compared two modeling algorithms (MaxEnt *vs* Random Forests), and two background point selection strategies (random points *vs* weighted points accounting for sampling effort). As an example application, we used SDM predictions to evaluate the conservation value of different protected area types within California.

**Results:** ClimateNA outperformed WorldClim for 94% of species, Random Forests outperformed MaxEnt for 87%, and random background points outperformed weighted background points for 100%. All differences were statistically significant. Together, the ClimateNA dataset, Random Forests, and random background points achieved highest performance for 86% of species. Using this best-performing set of models, we found that regional parks, county parks, state beaches, and open spaces in California were highest in multi-species suitability, while larger protected areas, such as national parks and national forests, generally exhibited surprisingly low suitability. Substantial spatial biases intrinsic to SDMing with unprojected predictor datasets (e.g., WGS84) are described, along with clear solutions using equal-area predictor datasets.

**Main conclusions:** Considerable disparity was observed among the performance of common SDM methods. This study highlights the importance of fine-scale, equal-area predictor datasets and best-practice guidelines, and demonstrates how SDMs can provide critical insights into protected area planning.

## Introduction

Understanding associations between species’ occurrences and variation in bioclimatic, ecological, and geographical variables is fundamental to many research fields, including ecology, evolution, biogeography, and conservation biology. Species distribution models (SDMs), sometimes called ecological niche or habitat suitability models, are powerful tools that quantify these associations and make spatially explicit predictions of where species are likely to occur (Elith and Leathwick, 2009a). These predictions are often used in real-world conservation decision-making, including management of at-risk species and the design of protected areas (Elith and Leathwick, 2009b; Wilson et al., 2011; Guisan et al., 2013; Whitehead et al., 2017). Because occurrence data are more readily available than true presence/absence data, presence-only modeling methods are widely applied in SDMing using algorithms such as MaxEnt (Phillips et al., 2006) and Random Forests (Breiman, 2001). As inputs, these models require only species occurrence records and bioclimatic, ecological, and geographical predictor variables; true absences are not required. Within the last decade, an explosion of community science species occurrence data, enabled by user-friendly resources like iNaturalist, has allowed SDMing greater taxonomic breadth than ever before (Beninde et al., 2023). Similarly, as fine-scale bioclimatic, ecological, and geographical data become increasingly available, it is becoming ever easier to generate diverse sets of predictor variables and produce ecologically meaningful models.

With little-to-no upfront cost, SDMing is becoming increasingly accessible to a diverse set of users, including both academic researchers and conservation managers. However, generating SDMs is a cumbersome process with an abundance of methodological recommendations (Araújo et al., 2019; Valavi et al., 2022). Because of the multitude of different methods, parameter settings, and available spatial data, model predictions are generally not comparable among studies and drawing real-world inferences can be complicated (Hallgren et al., 2019; Muscatello et al., 2021; Valavi et al., 2022). Best-practice guidelines for SDMing are valuable resources (see Araújo et al., 2019) for most users, leading to both better models and reasonable interpretation of their outputs and predictions. At the same time, there is a need to continually update these guidelines and to provide users with pre-processed, state-of-the-art bioclimatic, ecological, and geographical predictor datasets. These contributions streamline high-quality SDMing, adding to its accessibility in both research and applied conservation settings.

In this study, we generate and evaluate new bioclimatic, ecological, and geographical predictor variables using various approaches to SDMing, and, as an example application, use the resulting model predictions to assess the efficacy of different types of protected areas in California. Many high-resolution bioclimatic, ecological, and geographical datasets are already available for California. However, most of their spatial data layers do not extend far, or even at all, past the state’s borders. Examples include datasets available through USGS Basin Characterization Model (Flint et al., 2013) and the California State Geoportal (https://gis.data.ca.gov/). These spatial data have proven useful for landscape modeling at narrowly defined spatial extents, such as the California Floristic Province (Rose et al., 2023) and the Greater Los Angeles metropolitan area (Beninde et al., 2023). However, data layers restricted to state borders preclude some critical landscape modeling applications at broader spatial extents. For example, to accurately map species ranges and suitable habitats, predictor variables should be sampled from areas surrounding species presence points (i.e., beyond state borders), to allow theoretically available but unoccupied habitats to contribute to model training (Barve et al., 2011; Fourcade et al., 2014). Data layers restricted to state borders are also problematic for state- wide habitat connectivity analyses, which require all spatial data layers to fill a minimum convex polygon surrounding all points of interest (e.g., species occurrences used as nodes in circuit-based analyses). Ideally, this minimum convex polygon should be buffered by a large distance (e.g., >20% width of the study extent, or as a function of dispersal ability of species) to avoid underestimating landscape permeability at the edges of the modeling extent (Koen et al., 2014). To meet these requirements, all spatial data layers generated and evaluated in this study extend well beyond California’s state borders, allowing robust SDMs and connectivity models to be generated and used in California conservation initiatives.

California harbours substantial native and endemic biodiversity and is a recognized global biodiversity hotspot (Myers et al., 2000). It is also the most populous state in the US, harboring ∼12% of the country’s population in ∼5% of its continental land area. As a result, the state has many species of conservation concern, with more than twice as many federally protected species as any other mainland US state (292 at the time of writing; U.S. Fish and Wildlife Service, n.d.). Despite its high human population and associated ecological impacts (Knudson, 2005), California is also a model for conservation efforts, leading the USA in the maintenance and development of protected areas (Fiedler et al., 2022). California hopes to protect 30% of its lands and coastal waters by 2030, as envisioned by the state 30×30 initiative (Galloway, 2021). Habitat loss and degradation are among the biggest present-day threats to biodiversity worldwide (Brooks et al., 2002), meaning these habitat-based conservation initiatives are among the most important mechanisms for curbing biodiversity loss. Yet, the establishment of protected areas in the US and globally often prioritizes regions that are unsuitable for human activities and occupation, such as those at higher elevations, with steeper slopes, and at greater distances from urban centers, rather than areas that might yield the greatest conservation benefits (Nash, 2014; Joppa and Pfaff, 2009). In other words, protected areas are often allocated by default rather than by design, based on the best available scientific knowledge. Given its high biodiversity and prevalence of human activities, California is an ideal landscape for assessing the relevance of SDMing for conservation planning and management.

We focus our SDMing efforts on terrestrial species that are broadly representative of another California-led conservation initiative, the California Conservation Genomics Project (CCGP; Shaffer et al., 2022; Fiedler et al., 2022). The CCGP aims to generate whole-genome sequence data for roughly twenty thousand individuals belonging to 235 species to help manage California biodiversity in the face of climate change, habitat loss, and other human-mediated threats. These species were selected based on several criteria, including having a broad range and some or all of that range being of conservation concern or special scientific interest. Of these 235 species, 208 are terrestrial and include plants, fungi, invertebrates, and vertebrates that together span much of the geographical, environmental, and phylogenetic diversity found in the state (Shaffer et al., 2022; Fiedler et al., 2022; Toffelmier et al., 2022). These species are thus well-suited for evaluating the performance of different sets of predictor variables and SDMing methods. Resulting SDM predictions can also be used to evaluate the efficacy of California’s current network of protected areas, assessing which types of protected areas exhibit high mean suitability across many species of conservation concern and scientific interest.

To bolster SDMing resources, knowledge, and applications in California and beyond, the objectives of our study are threefold: 1) produce a highly accurate, fine-scale, equal-area projected predictor dataset for SDMing in California, encompassing a minimum convex polygon of the state buffered by 200 km; 2) generate over 5000 SDMs for 127 terrestrial species of conservation concern or other scientific interest in California, assessing differences in model performance between a) different bioclimatic predictor datasets (ClimateNA *vs* Worldclim), b) different modeling algorithms (MaxEnt *vs* Random Forests), and c) different modeling inputs (random *vs* weighted background/pseudo-absence points); 3) generate mean habitat suitability surfaces for all native species using the best-performing approach to SDMing (ClimateNA + Random Forests + random background points), and use them to assess how well different types of protected areas in California preserve multi-species suitable habitat. We also describe and discuss several substantial biases and concerns relating to SDMing with unprojected spatial data (e.g., WorldClim [Fick and Hijmans, 2017] and PRISM [Daly et al., 2008] data), and provide recommendations for avoiding these biases.

## Methods

### Study area and focal taxa

We defined our study extent as a minimum convex polygon encompassing California, buffered by 200 km. This ensured that a diversity of theoretically available but unoccupied habitats were sampled in the model training process, and that study extent edges will not interfere with any future state-level connectivity analyses, even for highly mobile species. As input data for our presence-only SDMs, we downloaded all research-grade iNaturalist occurrence records for all 208 terrestrial species included within the CCGP (downloaded from GBIF.org). We chose to use iNaturalist occurrence records to train our SDMs because: 1) they are all relatively recent, mostly from the last decade, and likely represent extant populations; 2) location uncertainty is explicitly and consistently quantified; 3) “research-grade” species identifications are verifiable and generally reliable; and 4) California has many iNaturalist users, providing a rich source of occurrence records. All varieties and subspecies were aggregated to species level for our analyses. From our downloaded iNaturalist data, we filtered occurrence records with location uncertainty exceeding 300 m, thinned them to one occurrence record per template raster cell (see below for raster information), and then removed species with less than 30 occurrence records from the dataset, resulting in a final set of 127 species (Supplementary Materials; Table S1). These included 48 plant species, 2 fungi species, 25 invertebrate species, and 52 vertebrate species. Eight species are not native to the study area, including two insects (*Ceratitis capitata* and *Homalodisca vitripennis*) and six plants (*Bromus tectorum, Phragmites australis, Carduus pycnocephalus, Centaurea solstitialis, Convolvulus arvensis,* and *Cynara cardunculus*). We included them in analyses of model performance and excluded them from analyses of protected area habitat suitability.

### Predictor variables

Predictor variables in our SDMs included a series of geographic information system (GIS) data layers. Unless otherwise specified, all spatial data manipulations were completed in R using the *terra* package (Hijmans et al., 2022). We generated two different sets of predictor datasets and assessed their performance difference in SDMing; one generated using ClimateNA software (Wang et al., 2016) and the other using WorldClim 2 (Fick and Hijmans, 2017). All layers in the ClimateNA dataset were generated in NAD 1983 California (Teale) Albers Equal Area Conic (EPSG:3310, “NAD83/California Albers”) at a spatial resolution of 300 m x 300 m.

This California-specific projection is optimized for area calculation and is recommended by the California Department of Fish and Wildlife (2022) for state-wide mapping. Our chosen resolution of 300 m represents a compromise between quantification of bioclimatic, ecological, and geographical heterogeneity at relatively fine spatial scales and the computational power required for fine-scale spatial analyses. The WorldClim dataset was generated in the native coordinate reference system (CRS; WGS84) and resolution of WorldClim 2 (30 arc-seconds). Due to this CRS, the actual resolution of the WorldClim dataset varies with latitude, equal to ∼926 x 926 m at the equator and decreasing poleward. The average east-west dimension of raster cells within our study area is ∼736 m, ranging from ∼797 m in the south to ∼669 m in the north. North-south dimensions are ∼926 m and do not vary.

### Elevation and terrain indices

As a basis for elevation and terrain variables in the ClimateNA dataset, we downloaded a tiled collection of Digital Elevation Model (DEM) rasters from the USGS 3D Elevation Program (3DEP) at 1 arc-second resolution (U.S. Geological Survey, n.d.). After merging tiles, we reprojected and resampled the raster to NAD83/California Albers at a resolution of 300 x 300 m and cropped the raster down to the study extent. This raster then served as the basic template for generation of all other predictor variables within the ClimateNA dataset. Elevation data were included within the WorldClim 2 download. Thus, to obtain a template raster for the WorldClim dataset, we simply cropped the WorldClim elevation raster to the study extent in WGS84.

Using each of the ClimateNA and WorldClim elevation rasters, we generated a series of terrain indices (“terrain” function, *terra* R package) following Wilson et al., (2007). These included: 1) terrain ruggedness index, estimated as the mean of absolute differences between the elevation of a focal cell and each of the eight surrounding cells; 2) topographic position index, estimated as the difference between the elevation of a focal cell and the mean of the 8 surrounding cells; and 3) terrain roughness index, estimated as the difference between the maximum and minimum elevation of a focal cell and the eight surrounding cells. We also estimated a heat load index using the R package *spatialEco* (Evans and Ram, 2021; following McCune and Keon, 2002), which predicts solar radiation as a function of the slope and aspect of each cell.

### Surface water

We downloaded river/stream and lake vector data for the study area from the Hydrosheds database (Lehner and Grill, 2013; Messager et al., 2016). These vector data were buffered by 100 m, intersected with the template rasters, and turned into binary raster layers (value of 1 if a raster cell contained water, 0 otherwise). This was performed separately for lotic and lentic data, to create separate raster layers, as some terrestrial species are associated with only one or the other (e.g., the CCGP species *Hetaerina americana*, lotic only; Grether et al., 2023). We generated a third raster for each dataset, measuring the shortest distance from each raster cell’s centroid to any surface water.

### Land cover

Land cover GIS data were acquired from the Commission for Environmental Cooperation (Commission for Environmental Cooperation, n.d.), generated using 2020 Landsat satellite imagery. The fine native resolution (30 m) of these data meant no downsampling was required when reprojecting and resampling (bilinear method) to match our template rasters. Fifteen land cover categories are present within the study area, including different types of human land use (e.g., “urban” and “cropland”; Commission for Environmental Cooperation, n.d.).

#### Urbanization

As a proxy for urbanization, we downloaded Nighttime Light maps from the Earth Observation Group’s Version 4 DMSP-OLS Nighttime Lights Time Series (Elvidge et al., 1997; Baugh et al., 2010; native resolution of 30 arc-seconds). From this dataset, we reprojected and resampled the average of the visible band digital number values to match template rasters. Values ranged from 0-63, with no missing data present within the study extent.

### Vegetation

To quantify the amount of photosynthetic activity on the landscape, we calculate the Enhanced Vegetation Index (EVI) (Jiang et al., 2008) using the entire MODIS AQUA (MYD09A1) (Vermote, 2021a) and TERRA (MOD09A1) (Vermote, 2021b) collection. We generated EVI raster layers for both the ClimateNA and WorldClim datasets using Google Earth Engine (Gorelick et al., 2017). Median EVI values from 2000 to 2023 were used. While the use of Normalized Difference Vegetation Index (NDVI) is more widespread, it is susceptible to error and uncertainty over various atmospheric and canopy background conditions. EVI adjusts for canopy background noise and atmospheric conditions while also having higher sensitivity in high biomass regions (Matsushita et al., 2007). We chose median values rather than mean because the annual distribution of EVI values are often moderately to highly skewed (Dong et al., 2019).

### Soil data

We downloaded a series of soil data layers from SOILGRIDS (ISRIC - World Soil Information, n.d.; Poggio et al., 2021), including organic carbon stocks (tons/ha), bulk density of the fine earth fraction (cg/cm³), and cation exchange capacity (mmol(c)/kg) for the first 30 cm of substrate. All missing data (<1% of study area) were interpolated using the average values of cells within 10 km (NA’s not considered). The native resolution of these layers is 250 m and required resampling and reprojecting to match the ClimateNA and WorldClim template rasters.

### Bioclimatic data: ClimateNA

We compiled data for 17 bioclimatic variables using ClimateNA v7.30 software (Wang et al., 2016), including mean temperature and precipitation of the summer, fall, winter, spring, and the entire year, the difference between mean temperatures of the warmest and coldest months (“continentality”), degree-days below 0°C (chilling degree days), degree-days above 5°C (growing degree days), mean annual precipitation as snow, mean annual relative humidity, and length of the frost-free period. ClimateNA generates scale-free bioclimatic data for user-specified locations using a combination of bilinear interpolation and local elevation adjustment and can be more accurate than traditional gridded data like PRISM (Daly et al., 2008) or WorldClim (Fick and Hijmans, 2017). Because these data are scale-free, no resampling/downsampling is required to match the template raster of a project’s analyses, which would result in decreased accuracy. We constructed 17 ClimateNA raster layers using our template raster cell’s centroid coordinates as inputs. However, only lat/long WGS84 coordinates (in csv file format) are accepted by the software. To build raster layers in an equal area projection (e.g., NAD83/California Albers) without reprojecting and resampling, we developed the following protocol: Working in NAD83/California Albers, we extracted all cell centroid coordinates from our template raster (populating columns 1 and 2 in a csv file). Each row in this file is a raster cell centroid. We then reprojected these centroid coordinates to lat/long WGS84 coordinates and populated columns 3 and 4 in the csv file and queried ClimateNA with these values. The resulting ClimateNA output file was identical in structure to the input file, and contained 17 new columns with the interpolated values of each bioclimatic variable. We then transformed these data into 17 raster layers (one layer per variable) using columns 1 and 2 in the csv file (NAD83/California Albers coordinates). Using this approach, we generated ClimateNA rasters at 300 x 300 m cell size, which is approximately 9X finer than most other climate data used for SDMing (e.g., WorldClim 2.1, Fick and Hijmans, 2017).

### Bioclimatic data: WorldClim

For comparison with our ClimateNA dataset, we downloaded all 19 bioclimatic WorldClim 2.1 variables (Fick and Hijmans, 2017) for the study area in their native CRS and resolution (WGS84, 30 arc-seconds). These 19 variables describe various aspects of temperature and precipitation patterns, including annual means, ranges, seasonal variations, and conditions during the wettest, driest, warmest, and coldest periods.

### Species distribution model (SDM) training, prediction, and evaluation

We built SDMs using two popular presence-only algorithms, MaxEnt (Phillips et al., 2006) and Random Forests (Breiman, 2001). These consistently rank among the best-performing and most computationally efficient approaches to correlative SDMing (Harrigan et al., 2014; Valavi et al., 2021). Using both MaxEnt and Random Forests, we compared: 1) our two bioclimatic predictor datasets (ClimateNA *vs* WorldClim); and 2) two different methods of generating background/pseudo-absence points (random *vs* weighted).

SDMs were trained using a subset of predictor variables selected to minimize collinearity (|*r*| < 0.7) and enable model transfer; i.e., make predictions across space or through time to different bioclimatic conditions (Guisan and Thuiller 2005; Elith and Leathwick 2009; Peterson et al., 2011). In all SDMs, our predictor variables included the terrain ruggedness index, the heat load index, land cover type, lotic water, lentic water, distance to any surface water, soil organic carbon, soil bulk density, soil cation exchange capacity, EVI median, mean temperature of the warmest quarter, mean temperature of the coldest quarter, difference between mean temperatures of the warmest and coldest months (ClimateNA) or quarters (WorldClim) (hereafter, “continentality”), mean precipitation of the warmest quarter, and mean precipitation of the coldest quarter.

In both ClimateNA and WorldClim SDMs, we compared the use of random *vs* weighted background/pseudo-absence points in model training. In the MaxEnt modeling process, the term ’background point’ is more commonly used because these points are used to characterize the range and distribution of predictor variables across the landscape. MaxEnt combines background points with occurrence records to sample the landscape and estimate the relative likelihood of species presence by contrasting occurrences with the background environment. Here, background points are not treated as absences. In contrast, Random Forests classification treats background points as pseudo-absences, assuming they represent areas where the species is not present. Random Forests aims to best separate pseudo-absences from occurrence records by identifying splits in the predictor variables that maximize classification accuracy. Hereafter, we use the term “background points” in all cases for simplicity.

Random background points are frequently used in SDMing. For example, they are often generated by default in the MaxEnt modeling process (Phillips et al., 2006). We generated our own to ensure comparability between models. As a basis for each species’s random background points, we first generated 50,000 points across the entire study extent and then cropped these points using a 200 km buffered minimum convex polygon surrounding that species’s occurrence records. This spatial constraint means that only environments that may be theoretically available to each species are sampled in the model training process. Further, failing to consistently constrain modeling extent can substantially inflate estimates of predictive power in model evaluation (Lobo et al., 2008; Sofaer et al., 2019). While random background points are most commonly used, this approach has been criticized for failing to account for spatial sampling biases that are present in many datasets, including iNaturalist (Kramer-Schadt, et al., 2013; Fithian et al., 2015; Barber et al., 2022; Beninde et al., 2023). As a solution, some have suggested using “weighted” background points that are spatially scaled in abundance in proportion to inferred sampling effort; i.e., generating background points with the same bias as occurrence data (Phillips et al., 2009; Kramer-Schadt et al., 2013; Searcy and Shaffer, 2016). To accomplish this, we downloaded research grade iNaturalist occurrence records for all plant and animal species within the study extent (GBIF.org, 2024a). We then randomly selected 50,000 plant occurrences and 50,000 animal occurrences for use as weighted background points in plant/fungi and animal SDMs, respectively. These were also cropped to the 200 km buffered minimum convex polygon surrounding each species’s occurrence records.

All SDMs were trained using the filtered and thinned dataset of iNaturalist occurrence records. For each species, we split occurrence records into five equal bins, each withholding a different 20% of records for model evaluation (five *k*-fold models). MaxEnt models were trained using the R package *dismo* (Hijmans et al., 2017). Following default settings, the regularization parameter was set to 1.0 and the number of feature classes was determined automatically based on the number of occurrences used (<10, only linear; 10-15, linear and quadratic; >15, linear, quadratic, and hinge; >80, all feature classes). Approximately 89% of MaxEnt SDMs included more than 80 occurrence records, meaning most MaxEnt SDMs used all feature classes. Random Forest SDMs were trained using the R package *randomForest* (Liaw and Wiener, 2022) using 500 trees and a “down-sampling” approach. This fitted each tree with a bootstrapped sample of occurrences and an equal number of randomly selected background points, which has been shown to drastically increase Random Forest SDM performance (Chen et al., 2004; Valavi et al., 2021).

### Prediction and evaluation

For each of the five *k*-fold models, we generated a predicted habitat suitability surface covering the entire study extent, with all raster cells receiving habitat suitability scores between 0 and 1 (higher values indicate higher suitability). MaxEnt model predictions were generated using the “clog-log” output and Random Forest predictions using “prob”. For each species, we summarized the five *k*-fold predicted habitat suitability surfaces using both the mean and coefficient of variation (CV) of each raster cell. Each species’ mean surface represents the final prediction of habitat suitability for a specific modeling method combination (e.g., ClimateNA + MaxEnt + random background points). The CV surface represents a direct index of prediction confidence–higher CV values indicate more variation among *k*-fold models and thus higher prediction uncertainty.

We evaluated the predictive power of all SDMs using area under the receiver operating characteristic curve (AUCROC). In all cases, AUCROC was completed using 50,000 random points generated across the entire study extent and then cropped to the extent at which each species’ SDMs were trained. For each combination of species, predictor dataset, modeling algorithm, and background point selection strategy, we calculated the mean AUCROC of all *k*-fold models to compare model fit across methods. AUCROC is frequently employed in SDM evaluation (Araújo et al., 2019; Valavi et al., 2022) and considers both the false positive rate (1 – specificity) and true positive rate (sensitivity) across all possible thresholds of predicted habitat suitability. Output values range from 0 - 1, with 0.5 indicating no discriminating power (uninformative prediction) and 1 representing perfect discrimination (Pearce and Ferrier, 2000).

For our best performing combination of SDM methods (ClimateNA + Random Forests + random background points), we also generated another set of SDMs for each species (five *k*-fold models) using the entire study extent instead of each species’ constrained extent. This ensured that the entire extent was sampled in the modeling process, allowing us to more accurately predict habitat suitability across the entire extent. Averaging the mean surface of all 119 native species (each scaled between 0 and 1) produced a mean multi-species suitability surface. To assess whether species “clustered” in their habitat associations, either taxonomically or functionally, we first extracted suitability scores for each species at 50,000 random points across the entire study extent and then completed a principal component analysis (PCA) on extracted values. Discrete clustering in PCA space would indicate that sets of species are similar in their habitat associations and different from other such sets of species. Conversely, no clustering would indicate that variation among species’ habitat associations is continuously distributed and not clustered. We also assessed the distribution of pairwise correlation coefficients among species’ predicted suitability surfaces. Multiple peaks in this distribution would indicate functional clusters of species, while a single peak would indicate that variation in species’ habitat associations is continuous.

### Methodological comparison

We used generalized linear mixed models (GLMMs) to compare performance among SDMs trained using: 1) different bioclimatic predictor datasets (ClimateNA *vs* WorldClim); 2) different modeling algorithms (MaxEnt *vs* Random Forests); and 3) different background point selection strategies (random *vs* weighted). Each of these modeling approaches was included in GLMMs as a binary variable, and all three were simultaneously regressed on AUCROC scores while controlling for non-independence among each species’s *k*-fold models including species ID as a random effect. We evaluated the effect of each binary variable using standardized coefficients. We also included the number of occurrence records used to train each SDM as a covariate to test for the effect of sample size on SDM performance. An interaction between modeling algorithm and number of occurrence records tests whether the effect of sample size on model performance differed between MaxEnt and Random Forests. We also reran GLMMs for all species that had fewer than 80 occurrence records in all their SDMs to assess whether the observed differences remained consistent at small sample sizes.

### Evaluation of California protected areas

We used the mean multi-species suitability surface (averaging all 119 native species’ suitability surfaces across the entire study extent) to evaluate how well different types of protected areas in California preserve multi-species suitable habitat. We downloaded shapefiles of all protected areas in California from the 30×30 Conserved Areas dataset (California Natural Resources Agency, 2023), which includes the California Protected Areas Database (CPAD) and California Conservation Easement Database (CCED). Polygons were combined by protected area name (e.g., “Yosemite National Park”) and then filtered for a minimum area of 1 km², resulting in the following sets of protected area types: 1) BLM lands (*n* = 897); 2) national forests (*n* = 20); 3) national parks (*n* = 9); 4) conservation easements (*n* = 1,418); 5) nonprofit lands (*n* = 1,071); 6) recreation areas (*n* = 64); 7) state parks (*n* = 83); 8) regional parks (*n* = 67); 9) open space (*n* = 100); 10) state beaches (*n* = 11); and 11) county parks (*n* = 21). We also collated polygons for all urban areas as defined by the U.S. Census Bureau (2020; *n* = 191). All polygons are non-overlapping, except conservation easements, nonprofit lands, and urban areas, which sometimes have multiple designations (e.g., a conservation easement may be managed by a nonprofit, and a state park may be included within an urban area). Mean suitability was extracted for all protected area and urban polygons and summarized by area type for comparison.

## Results

### Focal Taxa

After filtering iNaturalist occurrence records for accuracy, spatial proximity, and abundance, 127 terrestrial species remained for training SDMs, including 119 native species and eight non-native species. Plant species included Liliopsida (total *n* = 3; non-native *n* = 2), Magnoliopsida (*n* = 43; non-native *n* = 4), Marchantiopsida (*n* = 1), and Polypodiopsida (*n* = 1); both fungi species were in the Lecanoromycetes (*n* = 2); invertebrate species included Arachnida (*n* = 1), Diplopoda (*n* = 3), Gastropoda (*n* = 5), and Insecta (*n* = 16; non-native *n* = 2); and vertebrate species included Amphibia (*n* = 3), Aves (19), Mammalia (*n* = 11), Reptilia_Squamata (*n* = 18), and and Reptilia_Testudines (*n* = 1). Because occurrences were thinned by ClimateNA or WorldClim template raster cells (300 x 300 m and ∼736 x 926 m, respectively), training of SDMs using ClimateNA dataset supported the inclusion of ∼21% more occurrence records than the WorldClim dataset (Supplementary Materials; Fig. S1). In total, across all species, we used 219,228 occurrence records in ClimateNA models and 163,929 occurrence records in WorldClim models.

### Species Distribution Models

A total of 5080 SDMs were trained for 127 species. We observed considerable disparity in SDM performance in our comparisons of bioclimatic predictor datasets (ClimateNA *vs* WorldClim), modeling algorithms (MaxEnt *vs* Random Forests), and background point selection strategies (random *vs* weighted) (Fig. 1). Based on AUCROC scores, SDMs trained using the ClimateNA dataset outperformed those using the WorldClim dataset for 120/127 species (Supplementary Materials, Table S1), Random Forests outperformed MaxEnt for 111/127 species, and random background points outperformed weighted background points for all species. Overall, the best-performing SDMs by a substantial margin were those trained using the ClimateNA dataset, Random Forests, and random background points (109/127 species, including all 8 non-native species). For this set of models, mean predicted suitability across all 119 native species was generally highest around the coast and the Sierra Nevada mountain range (Fig. 2, panel a). Across the three analyses shown in Figure 2, we infer that: 1) species’ suitability scores do not average out to a uniform surface–there are consistent spatial patterns in habitat associations; 2) areas predicted to have higher mean multi-species suitability generally exhibited lower interspecific variation in suitability; and 3) considering species independently, we are also more confident in our predictions of high suitability than low suitability. No clustering was observed in our PCA analyses of species’ suitability surfaces (Supplementary Materials; Fig. S2). Pairwise correlation coefficients between native species’ predicted habitat suitability surfaces ranged from -0.578 to 0.982. Overall, species’ suitability surfaces were positively correlated (mean *r* = 0.400, s.d. = 0.341) and only a single peak was observed in a histogram of correlation coefficients (Supplementary Materials; Fig. S3).

**Figure 1.**
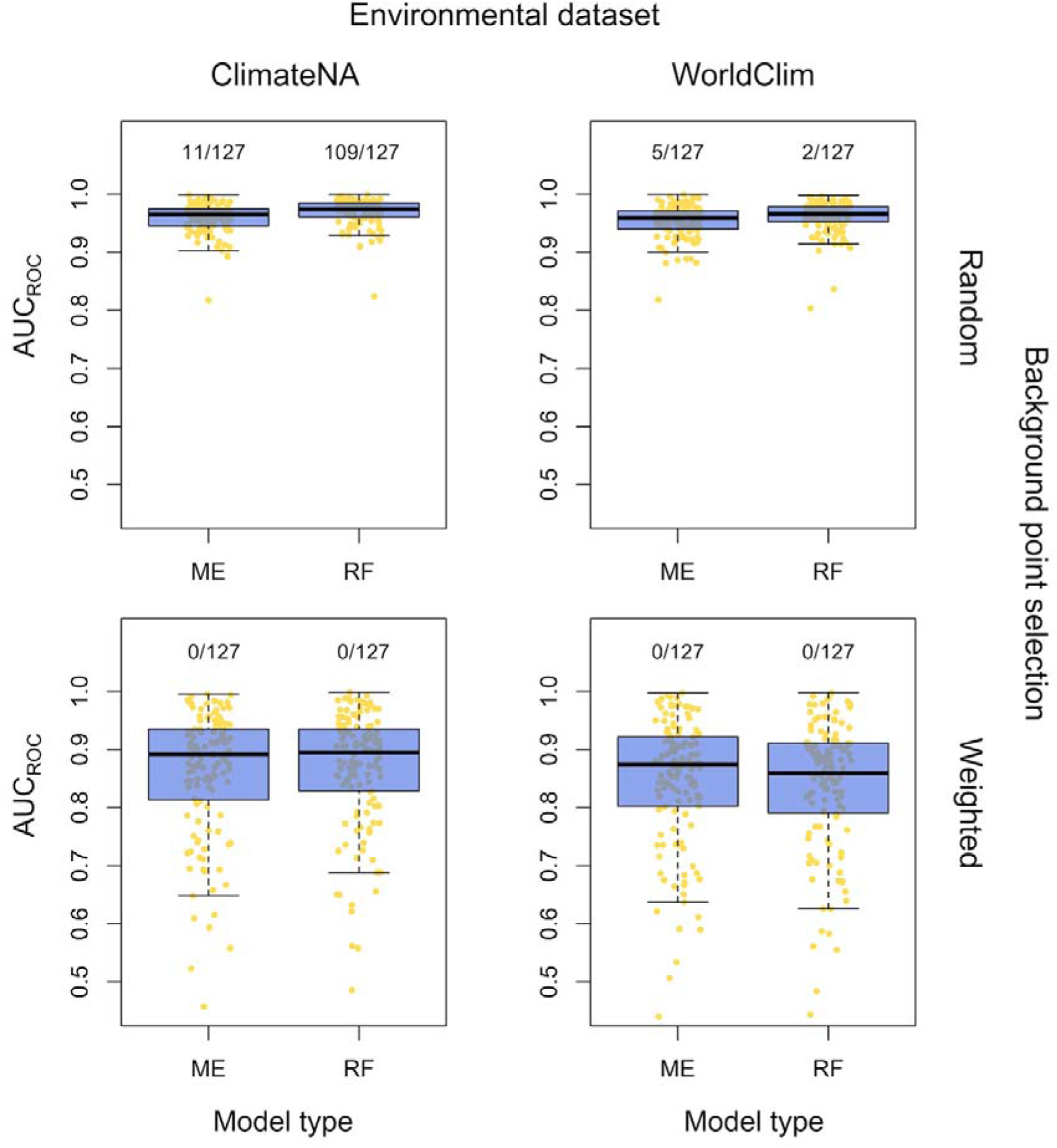
Species distribution model (SDM) performance, evaluated for 127 species using different predictor datasets (ClimateNA *vs* WorldClim), modeling algorithms (MaxEnt [ME] *vs* Random Forests [RF]), and background point selection strategies (random *vs* weighted). Within plots, each point represents the mean AUCROC score of the 5 *k*-fold SDMs for each species. The biggest difference in SDM performance was between background points (random *vs* weighted), followed by predictor dataset (ClimateNA *vs* WorldClim), followed by modeling algorithm (MaxEnt *vs* Random Forests). Boxes and whiskers superimposed on the points represent upper/lower quartiles and 95% confidence intervals, respectively. The fraction over each set of points represents the number of species for which the given model method combination was best performing. For example, for 109/127 species (including the eight non-native species), the best-performing SDM was built using the ClimateNA dataset, Random Forests, and random background points.

**Figure 2.**
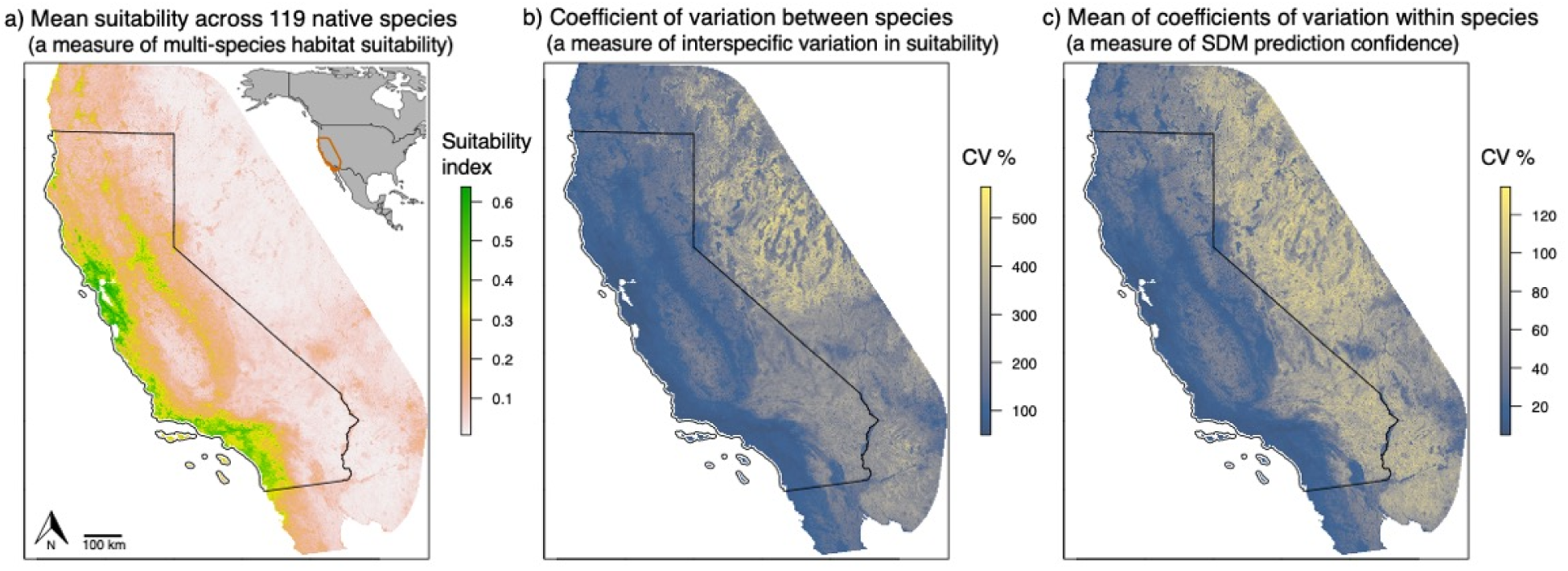
Predictions of species distribution models (SDMs) for 119 terrestrial species native to California and surrounding area, trained using our best-performing data and methods (ClimateNA dataset, Random Forests, random background points). a) Mean predicted suitability across 119 native species (the “mean multi-species suitability surface”) was generally highest around the coast and the Sierra Nevada mountain range. b) The coefficient of variation (CV) among species’ predicted suitability surfaces reflects the degree of interspecific variation in suitability within the study area (raster cells with a high CV indicate high interspecific variation in predicted suitability values). c) The CV among each species’ 5 *k*-fold models is proportional to the confidence we have in the suitability prediction (higher CV among predicted values equate to lower confidence in that region). The surface shown here represents the average CV values across the 119 species.

GLMMs confirmed that bioclimatic predictor dataset, modeling algorithm, and background point selection strategy all had significant effects on SDM performance (alpha = 0.05; Table 1). Based on observed effect sizes, the biggest difference in SDM performance was between background point selection strategies (random *vs* weighted), followed by predictor datasets (ClimateNA *vs* WorldClim), and then by modeling algorithms (MaxEnt *vs* Random Forests). The number of occurrence records used to train SDMs had a significant positive effect on SDM performance. The interaction between modeling algorithm and number of occurrence records was significantly negative, indicating that Random Forests benefited more from increasing numbers of occurrence records than MaxEnt. Specifically, the improvement in predictive power with additional occurrence records was greater for Random Forests than for MaxEnt, suggesting that Random Forests is more sensitive to sample size increases. A total of 13 species had fewer than 80 occurrence records in SDMs. Rerunning GLMMs on the SDM performance for just these species shifted the results, suggesting that MaxEnt might outperform Random Forests when few occurrence records are available (Table S2). The directionality of other effects remained consistent, but due to the small sample size most were not significant. Raising the threshold for occurrence values to 100 and 120 reduced the significance of the algorithm effect, suggesting that once the number of occurrence records exceeds ∼80, Random Forests performs better on average. Lowering the threshold also reduced significance, likely due to the small number of species included in the regression. Further analyses (e.g., subsetting occurrences of all species) are required to actually resolve the number of occurrences at which relative algorithm performance changes.

**Table 1.** Generalized linear mixed models (GLMMs) compared performance among SDMs built using: 1) different bioclimatic predictor datasets (ClimateNA *vs* WorldClim); 2) different modeling algorithms (MaxEnt *vs* Random Forests); and 3) different background point selection strategies (random *vs* weighted background points). The best performing modeling methods (ClimateNA, Random Forests, and random background points) were used as reference categories for measuring effects. After thinning to one occurrence record per raster cell, training SDMs using the ClimateNA dataset included ∼21% more occurrence records than the WorldClim dataset given its finer resolution. We controlled for the effect of number of occurrences as a covariate, and tested whether its effect varied between modeling methods (MaxEnt *vs* Random Forests) using an interaction term. AUCROC scores were used as the response variable and non-independence among each species’s *k*-fold models was controlled for using species ID as a random effect.

| Effect | Estimate | Std. error | z-value | p-value |
| --- | --- | --- | --- | --- |
| (intercept) | 3.52 | 0.0789 | 44.7 | <0.001 |
| predictor dataset (WorldClim) | -0.155 | 0.0106 | -14.6 | <0.001 |
| modeling algorithm (MaxEnt) | -0.0596 | 0.0114 | -5.21 | <0.001 |
| background points (weighted) | -1.49 | 0.0116 | -129 | <0.001 |
| number of occurrences | 3.56E-05 | 8.45E-06 | 4.17 | <0.001 |
| modeling algorithm:number of occurrences | -7.89E-06 | 3.39E-06 | -2.33 | 0.0198 |

### Evaluation of California protected areas

We discovered considerable disparity in mean multi-species suitability among different types of protected areas in California (Fig. 3). Overall, county parks, open spaces, regional parks, and state beaches had the highest mean suitability. Urban areas also exhibited high multi-species suitability. Lowest suitability was observed within national parks, national forests, and BLM lands. Other protected areas exhibited intermediate mean suitability values. Considering all types of protected areas simultaneously, we observed a significant, negative relationship between protected area size and mean habitat suitability averaged across all 119 native species (log-transformed area: *r* = -0.317 p < 0.001). This relationship remained significant and negative when excluding BLM, nonprofit, and Easement areas (log-transformed area: *r* = -0.526 p < 0.001; Supplementary Materials, Fig. S4). These latter areas contained a substantially higher number of polygons in the analysis due to the lack of a common identifier for polygons belonging to the same administrative unit, which prevented merging of multiple smaller polygons into a single larger one.

**Figure 3.**
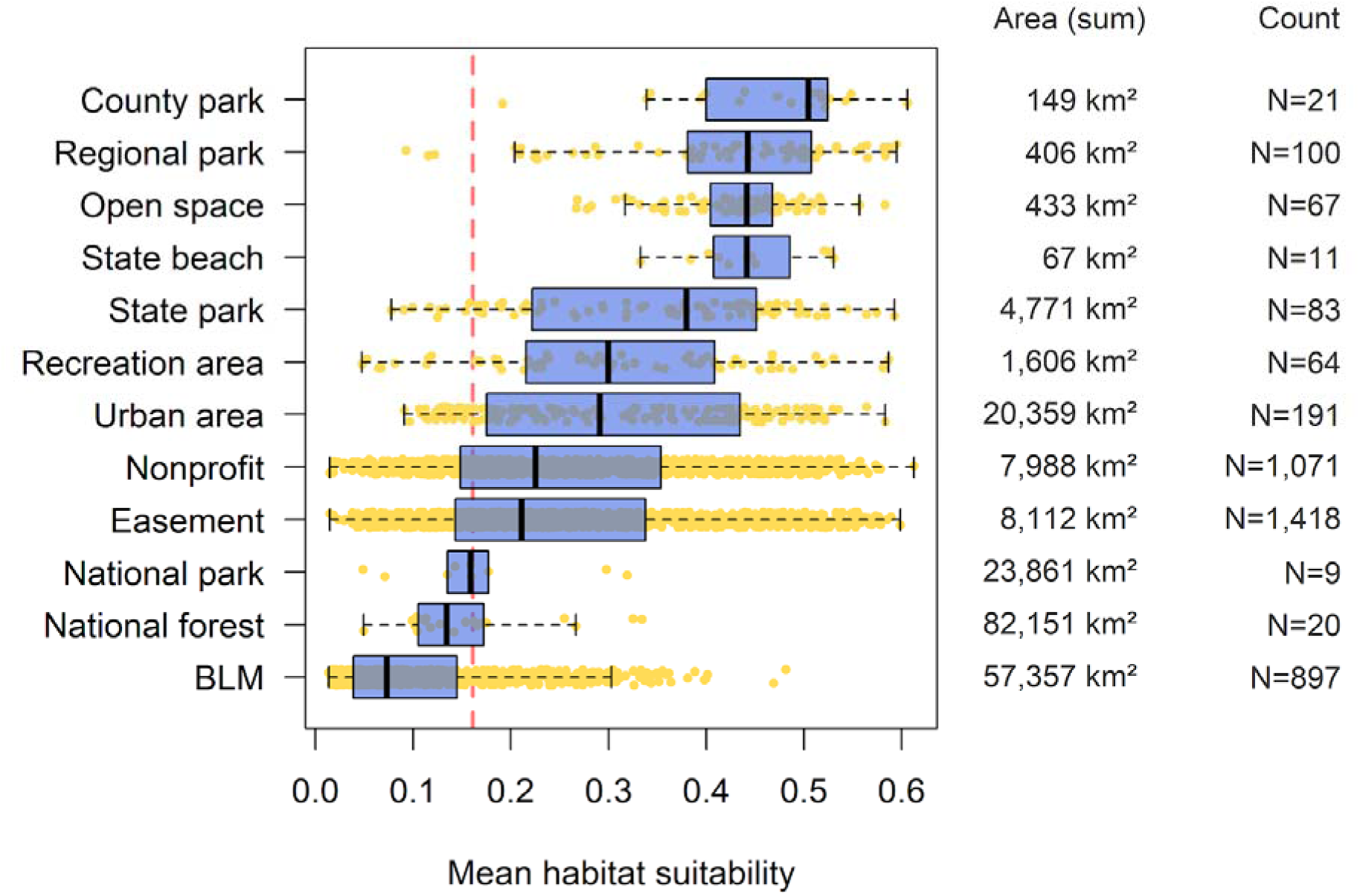
Mean multi-species suitability (119 native species) for protected and urban areas in California. Habitat suitability surfaces were generated for each species using the ClimateNA dataset, Random Forests, and random background points. Boxes and whiskers superimposed on points represent upper/lower quartiles and 95% confidence intervals, respectively. The dashed vertical line represents the mean multi-species suitability score for California.

## Discussion

Our study provides foundational resources for future SDMing in California: a new, fine-scale bioclimatic predictor dataset and best-practice SDM guidelines. We also provide reproducible code, allowing users to use these layers to generate SDMs and predicted suitability surfaces for any species of interest in California and surrounding area. Analyzing our 5080 SDMs in a comparative framework, we found that model training using our ClimateNA dataset, Random Forests algorithm, and random background points most effectively predicted suitable habitat for the vast majority (86%) of 127 terrestrial species of conservation concern or scientific interest. Given these species selection criteria (Shaffer et al. 2022), the average of all native species’ suitability surfaces (“mean multi-species suitability surface”) is arguably well-suited for broad-sweeping and preliminary evaluations of the conservation value of landscapes. Other studies have demonstrated significant associations of similar mean multi-species habitat suitability surfaces and species richness estimates derived from independent monitoring efforts, including plants in the Swiss alps (Brun et al., 2024) and butterflies in Italy (Riva et al., 2023). In California, our analyses show that multi-species suitability varies widely among federal, state, and nonprofit protected areas, highlighting that not all protected areas should be treated as equivalent in conservation planning and area-based conservation goals.

### Predictor dataset

A major product of this study is an open-access dataset of fine-scale bioclimatic, ecological, and geographical predictor variables specifically designed for terrestrial SDMing within California and surrounding area. This dataset includes what we have judged to be the most accurate and comprehensive elevation, terrain, water, land cover, urbanization, vegetation, and soil spatial data available for the study area, plus 17 bioclimatic variables generated using ClimateNA software. ClimateNA data have been shown, on average, to be more accurate than other bioclimatic data typically used in SDMing, such as WorldClim and PRISM (Wang et al., 2016). In topographically complex areas, bioclimatic conditions can vary at spatial scales much finer than 30 arc-seconds, the finest resolution available for WorldClim and PRISM (Wang et al., 2016). ClimateNA’s bilinear interpolation transforms these coarse baseline data into a seamless surface while using elevation adjustments to improve prediction accuracy for any specific locations of interest (Hamann and Wang, 2005; Wang et al., 2006). By specifying raster centroids as locations of interest (in WGS84 coordinates), ClimateNA allows users to generate highly accurate bioclimatic data layers for any spatial resolution. What we provide here is a simple method for generating data layers in an equal-area projection (rather than WGS84) without reprojecting/resampling, which can drastically reduce accuracy.

### Coordinate reference system (CRS)

It is important to recognize that any SDM built using WorldClim or PRISM data without reprojecting/resampling (which decreases data accuracy through an averaging process) will produce predicted habitat suitability surfaces with raster cells that vary in both area and shape across the study extent. WorldClim or PRISM data are generated and available in WGS84, meaning raster cells are defined in angular units (latitude and longitude) rather than true area-based units, leading to non-uniform spatial resolution because lines of longitude converge toward the poles. Within our study extent alone, 30 arc-second WGS84 raster cells vary in area from ∼0.737 km² in the south to ∼0.621 km² in the north, about 17% ([[largest cell area−smallest cell area]/average cell area]*100%). If occurrence records are filtered to one occurrence per raster cell, which is often the default practice (e.g., Phillips et al., 2006 and this study), variation in cell area will lead to variation in the density of occurrences across the study extent. This introduces important spatial bias in model training, incorrectly inferring more suitable conditions in poleward areas of the study extent than if occurrences were filtered uniformly. The magnitude of this bias will increase with increasing latitudinal range of the modeling extent. In some studies addressing small spatial extents, variation in raster cell area due to unprojected spatial data is small; e.g., the Greater Los Angeles Area has variation ∼1% (Beninde et al., 2023). In these situations, modeling using unprojected spatial data may be preferable over reductions in accuracy associated with resampling/reprojecting. However, because it is now possible to generate bioclimatic datasets in appropriate equal-area projections using ClimateNA and the methods described here, we recommend using them.

Spatial variation in raster cell area is similarly problematic for interpreting habitat area, configuration, and connectivity. When visually interpreting suitability surfaces that are not equal-area projected (e.g., WGS84), poleward areas will appear relatively larger, possibly biasing conservation managers’ interpretations of the distribution of suitable habitat. To correctly produce easily-interpretable suitability surfaces, users need to generate SDMs and predictions in equal-area projections optimized for their study area. For example, NAD83/California Albers is recommended by the California Department of Fish and Wildlife (2022) for state-wide, equal-area mapping. These considerations are particularly important for analyses of landscape connectivity, which are also one of the main goals of the CCGP (Shaffer et al., 2022; Fiedler et al., 2022). Predicted suitability surfaces are often used as inputs for resistance-based connectivity analyses, for example using Circuitscape and Omniscape (McRae et al., 2008; 2016), and implicitly assume uniform raster cell sizes when calculating cumulative resistance between locations on a landscape. To illustrate the magnitude of the bias introduced when using unprojected WGS84 raster surfaces for our study extent (California buffered by 200 km), we note that pairs of points at equal latitude in the extreme north of the extent (cell width = ∼669 m) will be estimated to be up to 19% more isolated than equidistant pairs in the extreme south of the extent (cell width = ∼798 m), introducing substantial spatial bias. This bias can be greatly reduced by projecting spatial data. Although not equal-area *per se*, Universal Transverse Mercator (UTM)-based projections are sometimes used in connectivity analyses (e.g., MacDonald et al., 2020 and some datasets analyzed by Beninde et al., 2024). One advantage of UTM-based projections is minimization of spatial distortion, i.e., the preservation of angles and shapes. Therefore, UTM-based projections may best preserve the relative configurations of nodes while minimizing biases associated with variation in raster cell area. However, this is only effective over relatively small spatial extents, such as single UTM zones, which are each 6 degrees of longitude wide. For each application, connectivity modelers should explore a variety of projections and report their selection.

Across SDMing and connectivity studies, the importance of these spatial biases is seldomly adequately addressed, and may even be entirely overlooked. More often than not, CRSs are not reported, or reported to be a non-equal-area projection, including SDM best-practice guidelines (Phillips, 2005; Phillips et al., 2006; Araújo et al., 2019), resistance-based connectivity modeling literature (e.g., McRae and Beier 2007; Pless et al., 2021; Vanhove and Launey, 2023; Beninde et al., 2024), and documentation of connectivity software (e.g., Circuitscape and Omniscape; McRae et al., 2008; 2016). A particular concern is that users will often adopt the default spatial definitions of data sources, which, as is the case for the widely-used WorldClim data, are frequently set to unprojected WGS84. The methods and bioclimatic predictor dataset provided in this study enable future SDMers to consistently and accurately produce equal-area predicted suitability surfaces, facilitating meaningful quantifications of habitat area, configuration, and connectivity.

### Resolution

The fine spatial grain of our predictor variables improves SDM performance, not only due to the increased accuracy of the spatial data, but also because it allows inclusion of a greater number of occurrence records for densely sampled taxa. Occurrences were thinned to one occurrence per ClimateNA or WorldClim template raster cell (300 x 300 m and ∼736 x 926 m, respectively), meaning training of SDMs using the ClimateNA dataset allowed inclusion of ∼21% more occurrence records than the WorldClim dataset in this study. Indeed, a principal benefit of high-resolution spatial data is the inclusion of a greater number of occurrence records. However, even though all occurrences included in training and testing data occur in different raster cells, this doesn’t mean the occurrences are truly independent (Bahn and McGill, 2012; Valavi et al., 2022). Lack of independence among clustered occurrences may inflate the estimated performance of SDMs trained using ClimateNA. Notwithstanding, our GLMMs showed that, even after controlling for the number of occurrence records, SDMs trained using the ClimateNA dataset still performed the best. We therefore recommend using this dataset for future SDMing efforts with California and surrounding area.

### Variable selection

In this study, we selected five bioclimatic variables to minimize collinearity. However, future users may choose to select other combinations of bioclimatic variables that are more appropriate for their species and modeling extent (collinearity will change across different modelling extents). Combining bioclimatic variables into uncorrelated composite variables via PCA is also possible (e.g., MacDonald et al., 2022). However, if users are only use SDMs to make predictions into the spatial extent and time period in which models were trained (i.e., not predicting across extents or into different time periods), collinearity may not be much of an issue for prediction accuracy (Guisan and Thuiller, 2005; Elith and Leathwick, 2009a; Peterson et al., 2011). Still, collinearity can mask the true importance of individual variables by distributing importance across correlated predictors, complicating model interpretation. In the Supplementary Materials we provide 17 bioclimatic variables generated via ClimateNA for future user selection.

### Modeling methods

#### Algorithm

Across 5080 SDMs, we observed that Random Forests significantly outperformed MaxEnt, providing the best prediction for 87% of species. This difference is best attributed to Random Forests’ flexibility, ability to capture complex interactions, and robustness to overfitting. Random Forests is a non-parametric machine learning algorithm that uses an ensemble of decision trees (Breiman, 2001). Through this process, it is also able to detect complex, nonlinear interactions between these variables without explicit feature engineering. Although MaxEnt can also model complex relationships using feature classes (linear, quadratic, product, threshold, and hinge), these transformations are predefined and may not fit highly nonlinear or complex relationships (Phillips et al., 2006). In this process, MaxEnt uses regularization to control overfitting, but, contingent on the correlation structure of predictor variables and the number of occurrence records used, models can become over- or under-fitted depending on the regularization parameter. Because of the large number of species and modeling approaches addressed here, we did not experiment with changing either feature classes or the regularization parameter. In contrast, Random Forests uses bootstrapping and random feature selection for each tree split to avoid overfitting. This is automatic and not contingent on sample size. For modeling of single or a few species, users may be able to increase the predictive power of MaxEnt SDMs by optimizing feature classes and the regularization parameter; e.g., using the R package *ENMeval* (Muscarella et al., 2014). This optimization is not required for Random Forests, streamlining the SDMing process and generally providing better results.

The only exception to our recommendation of Random Forests over MaxEnt is when modeling species with very few occurrence records. We observed that MaxEnt may outperform Random Forests for species modelled using less than ∼80 occurrence records. Although the effect size was observed to be small, this finding aligns with previous studies, which have demonstrated that MaxEnt performs better or comparably to other methods under sparse data conditions (Elith et al., 2006; Wisz et al., 2008). Our results suggest that MaxEnt’s ability to efficiently use background data and apply regularization to prevent overfitting is effective even when few occurrences are used. Thus, for small sample sizes, users may wish to experiment with both modeling algorithms.

#### Background points

Presence-only data, like iNaturalist, often have spatial biases due to uneven sampling effort. For example, areas near roads or urban centers may be oversampled, while remote areas may be undersampled, even though they may contain suitable habitat. This has led some SDMers to infer that accounting for sampling biases using weighted background points will improve SDM predictions (Kramer-Schadt et al., 2013; Beninde et al., 2023). If sampling bias is ignored, SDMs may incorrectly associate species’ occurrences with intensely sampled regions rather than true habitat suitability (Phillips et al., 2009, Fithian et al., 2015). However, our comparisons of background point selection strategies suggest that training SDMs using weighted background points that mirror sampling effort does not sufficiently sample the range of bioclimatic, ecological, or geographical variables present within the modeling extent. Using weighted background points significantly decreased SDM performance to a greater extent than any other variable (and all other variables combined). Other studies have reached similar conclusions (Beninde et al., 2023), and users can consult Valavi et al. (2022) for an empirical method for determining the optimal number of random background points.

It is important to note that our training and testing datasets were drawn from the same pool of occurrence records, meaning they are subject to identical sampling biases. We therefore cannot infer whether we modeled only species’ true habitat associations, or a combination of species’ true habitat associations and iNaturalist survey effort. Undoubtedly, using weighted background points produced poorly performing SDMs in this study. However, we cannot be sure the extent to which weighted background points actually controlled for sampling bias in the model training process, separating species’ true habitat associations and iNaturalist survey effort. To adequately assess this, model evaluation using independent datasets that do not contain spatial sampling biases are required. This was beyond the scope of this study.

### Conservation

The 119 native terrestrial species modeled here were selected by the CCGP on the basis of conservation concern and scientific interest (Shaffer et al., 2022; Fiedler et al., 2022). The taxonomic breadth of this initiative is broad, and our filtered dataset includes 48 plant, 2 fungus, 25 invertebrate, and 52 vertebrate species. These analyses heavily favor vertebrates and other large species, highlighting a need for increased research attention on invertebrates and other less-known groups (Toffelmier et al., 2022). Further, this selection of species seems to have favored those that occur along the coast and in human-occupied landscapes, with few mountain or desert specialist species; specialists were generally avoided in CCGP species selection due to their relatively confined ranges. Comparisons of predicted suitability surfaces showed considerable interspecific variability, with pairwise correlation coefficients between species’ surfaces ranging from -0.578 to 0.982. Still, we found no clustering in species’ habitat associations, suggesting this variation was continuously distributed. Overall, native species’ suitability surfaces were positively correlated (mean *r* = 0.405), and the mean multi-species suitability surface exhibited substantial variability across different types of protected areas in California (Fig. 3). Larger protected areas in California, such as National Parks and National Forest areas, had relatively low multi-species habitat suitability, while smaller protected areas, such as county parks, regional parks, and recreation areas, exhibit very high multi-species suitability. Urban areas also exhibited high multi-species suitability, emphasizing the intense competition for space between humans and native biodiversity and highlighting a need for strategic protected area planning. Globally, it is clear that protected areas are often established in places that are unlikely to be threatened by human development, even in the absence of protection (Joppa and Pfaff, 2009). California appears to be no exception, with most of its large, flagship protected areas established in areas that are not ideal for human occupation, following the “conservation by default” paradigm. However, other protected area types, such as state protected areas and parcels purchased and managed by nonprofit organizations, appear to be more suitable across many species, following the “conservation by design” paradigm.

The mean multi-species suitability surface generated here suggests California’s largest protected areas are, on average, of lower conservation value than other protected area types. Yet, BLM lands, national parks, and national forests frequently protect charismatic and expansive landscapes (e.g., Yosemite National Park) and sometimes protect unique, endemic species from specific threats (e.g., protecting the Desert Tortoise [*Gopherus agassizii*] from solar developments in Mojave National Preserve). Still, in our analyses, it is clear that not all protected areas are “equal” in multi-species conservation value, which deserves consideration in conservation planning. California’s 30×30 initiative, for example, aims to protect 30% of California’s land and coastline by 2030. This is an area-based goal that, depending on how it is implemented, may be agnostic to conservation quality. We hope that the SDMs and data products provided here will contribute to future protected area planning within the state and beyond, enabling prioritization of land based on differential conservation values. Future efforts could also explore spatial prioritization tools and data sets that explicitly account for the uniqueness of species, genetic resiliency of populations (Shaffer et al., 2022), and other intrinsic properties of species and populations, thereby allowing managers to set species-specific area- goals for species long-term persistence (e.g., Delavenne et al., 2012; Lessmann et al., 2014). In all of these future efforts, it would be best to explicitly investigate the effect of sampling biases on suitability surfaces, preferably using occurrence records from systematic surveys in which effort is standardized across space.

## Conclusions

Results of this study show that new methods for generating fine-scale predictor datasets, in combination with the Random Forests algorithm and random background point selection strategy, can generate SDMs and predicted suitability surfaces with a high degree of accuracy for a large diversity of terrestrial species in California. Without much fine-tuning, future users will be able to create similar high-quality SDMs and predicted suitability surfaces for any terrestrial species of interest within the study extent using the code provided here (Supplementary Materials). However, for projects focusing on single or small numbers of species, we recommend experimenting with different combinations of predictor variables and modeling methods. Applications of SDMs generated here include mapping suitable habitat, evaluating and planning protected areas, and inferring habitat connectivity. An important future research direction will involve using ClimateNA to generate a plethora of new bioclimatic datasets based on various climate change scenarios (Mahony et al., 2021). Using those data and the SDMs generated here, it will be possible to predict how the amount and configuration of suitable habitat is likely to change for terrestrial species of conservation concern and scientific interest, further informing protected area planning and California conservation science. Further, all methods presented here for generating predictor datasets can be similarly applied across all of North America, including Mexico, allowing users to generate SDMs for all terrestrial North American species with sufficient occurrence data.

## Contributions

Spatial data compilation and generation were completed by ZGM with assistance from JB and BZ. Species distribution models and suitability predictions were completed by KM with supervision from ZGM and JB. ZGM and JB completed subsequent analyses and made figures. JB compiled reproducible code. Writing was led by ZGM with assistance from JB and input from all authors.

## Acknowledgements

We thank Tongli Wang for assistance generating the ClimateNA data used in our analyses, Henry Chang (UCLA IT Support Center) for assistance with cloud storage, and members of the Shaffer Lab and California Conservation Genomics Project team for conceptual input.

