## Supplementary Materials for "Species distribution modeling for conservation science: new predictor layers, reproducible code, and an evaluation of California protected areas"

Table S1. 127 species are included in our species distribution modeling (SDMing) efforts. Listed are the best performing modeling methods for each species (including predictor dataset, model algorithm, and background point selection strategy), the total number of occurrence records per species after filtering to one record per ClimateNA raster cell, and the mean AUC_ROC_ score for the top performing combination of model methods. (Available as a downloadable .xlsx file.)

Table S2. Generalized linear mixed models (GLMMs) were used to compare performance among SDMs for species with less than 80 occurrence records after thinning using both the ClimateNA and WorldClim template rasters (*n* = 13). SDMs were built using: 1) different sets of bioclimatic predictors (ClimateNA *vs* WorldClim); 2) different modeling algorithms (MaxEnt *vs* Random Forests); and 3) different background point strategies (random *vs* weighted background points). Unlike GLMMs addressing SDMs for all species, MaxEnt was observed to significantly outperform Random Forests when sample sizes are very small. AUC_ROC_ scores were used as the responding variable and non-independence among each species’s *k*-fold models was controlled for using species ID as a random effect.

| Effect | Estimate | Std. error | z-value | p-value |
| --- | --- | --- | --- | --- |
| (intercept) | 3.11 | 0.62 | 5.02 | <0.001 |
| predictor dataset (WorldClim) | -0.0591 | 0.0704 | -0.84 | 0.401 |
| modeling algorithm (MaxEnt) | 0.423 | 0.204 | 2.07 | 0.0385 |
| background points (weighted) | -1.04 | 0.052 | -20.1 | <0.001 |
| number of occurrences | 0.00281 | 0.0101 | 0.279 | 0.78 |
| modeling algorithm:number of occurrences | -0.00744 | 0.00374 | -1.99 | 0.0465 |

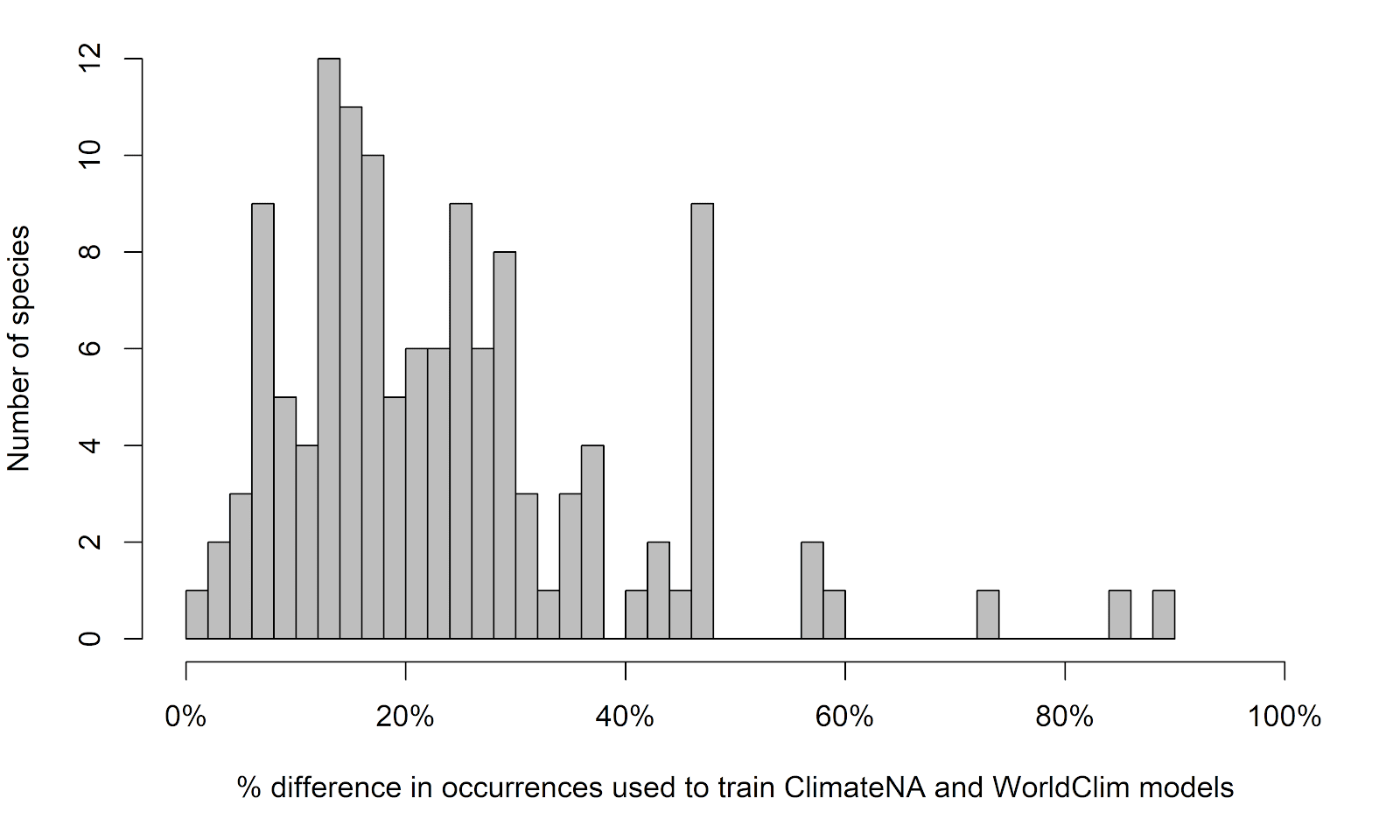

Figure S1: Histogram showing how many more occurrence records per species became available for SDMing when using ClimateNA over WorldClim due to the finer resolution data ([[N ClimateNA occurrences−N WorldClim occurrences]/N WorldClim occurrences]*100%).

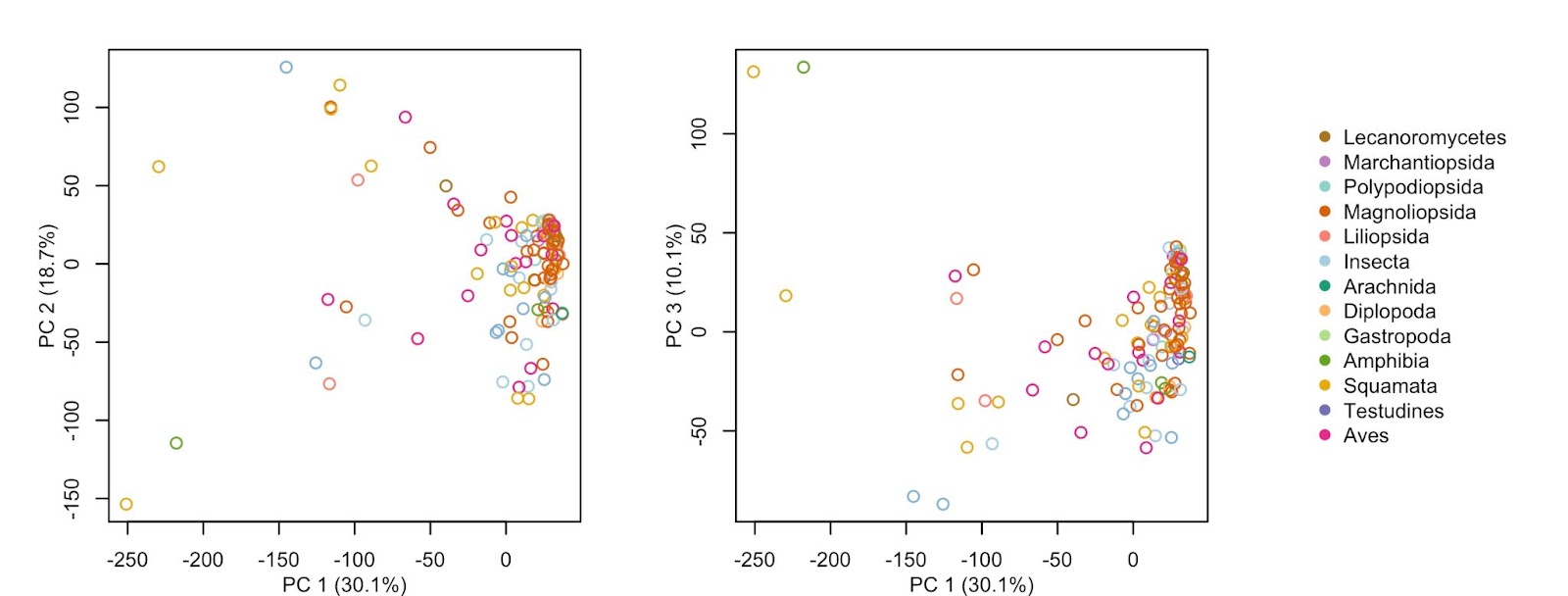

Figure S2. To assess similarity in species’ predicted habitat suitability surfaces, we extracted suitability scores for each species’s surface at 50,000 random points across the entire study area. We then ran a principal component analysis (PCA) on extracted values to assess whether particular groups of species exhibited clustering in PCA space or if variation among species was continuously distributed. No discrete clustering was observed.

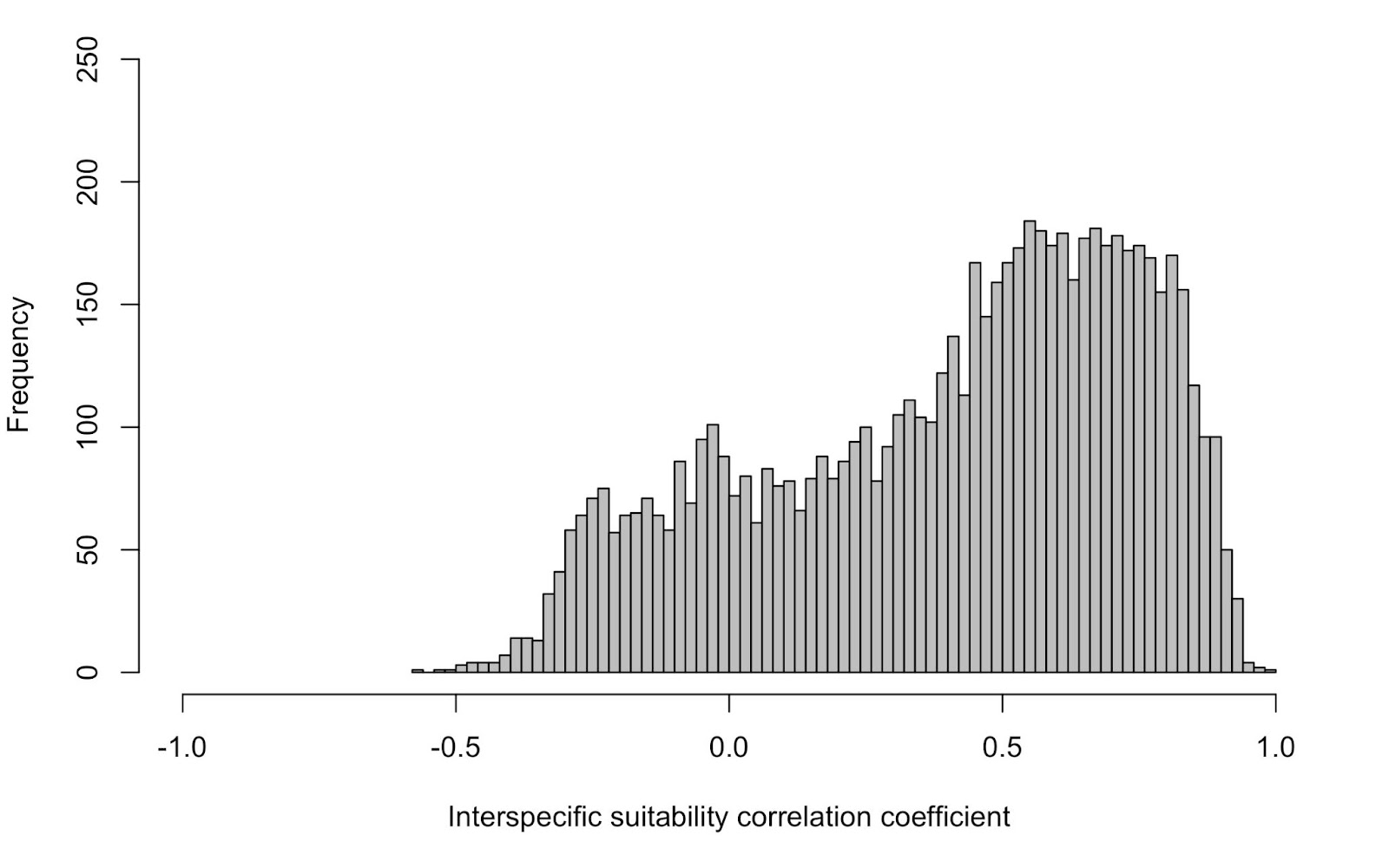

Figure S3. Histogram showing pairwise correlation coefficients between 119 native species’ predicted habitat suitability surfaces. Overall, species’ suitability surfaces were positively correlated (mean *r* = 0.400, s.d. = 0.341).

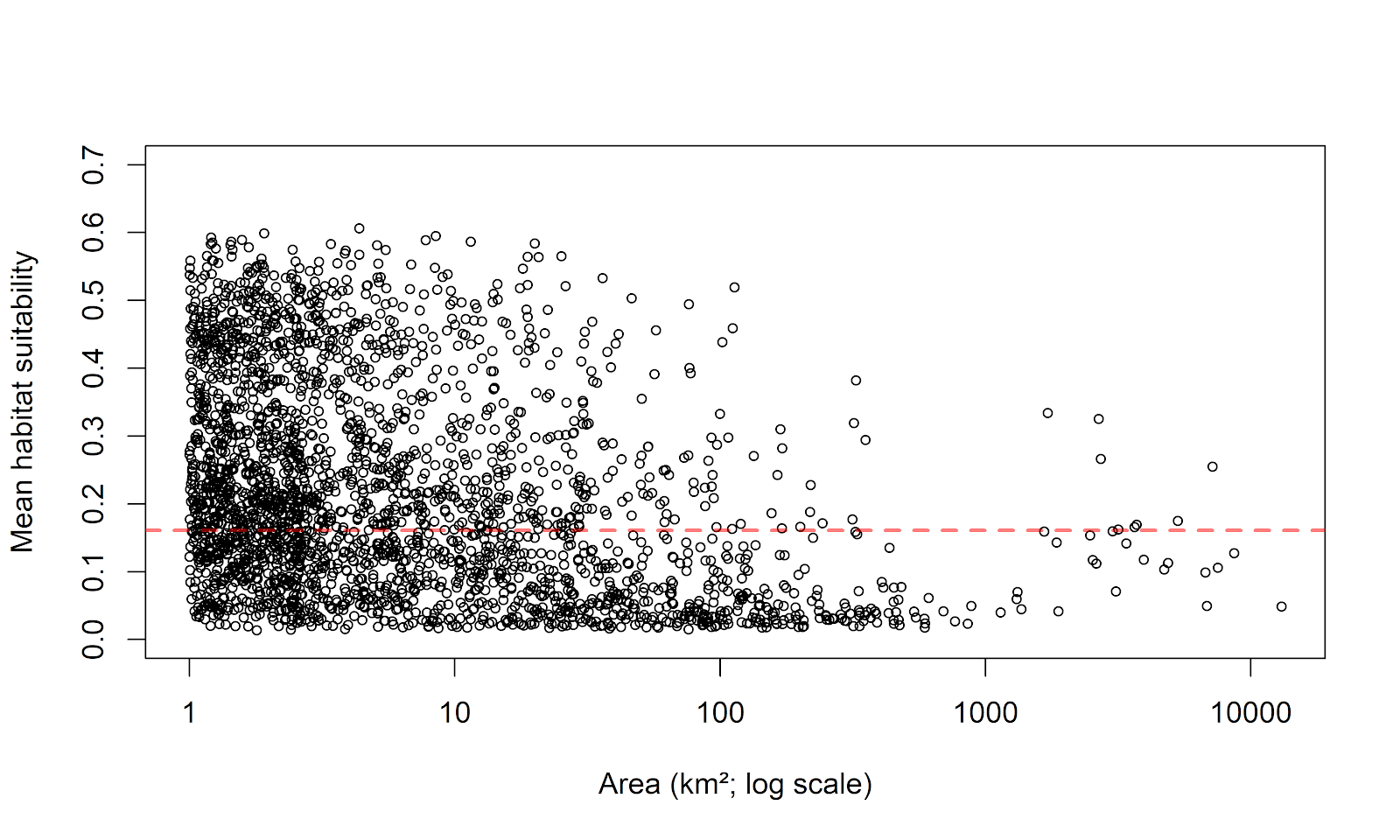

Figure S4. Relationship between multi-species suitability (averaged across 119 native species) and protected area size. All protected areas in California over 1 km^2^ were included. Pearson correlations were as follows: suitability of all protected areas and log-transformed area: *r* = -0.345 p < 0.001 (shown here); suitability of protected areas but excluding BLM, nonprofit, and easement areas and log-transformed area: *r* = -0.504 p < 0.001. The dashed horizontal line represents the mean multi-species suitability score for California.
